# AlphaTracker: A Multi-Animal Tracking and Behavioral Analysis Tool

**DOI:** 10.1101/2020.12.04.405159

**Authors:** Zexin Chen, Ruihan Zhang, Yu Eva Zhang, Haowen Zhou, Hao-Shu Fang, Rachel R. Rock, Aneesh Bal, Nancy Padilla-Coreano, Laurel Keyes, Kay M. Tye, Cewu Lu

## Abstract

The advancement of behavioral analysis in neuroscience has been aided by the development of computational tools^1,2^. Specifically, computer vision algorithms have emerged as a powerful tool to elevate behavioral research^3,4^. Yet fully automatic analysis of social behavior remains challenging in two ways. First, existing tools to track and analyze behavior often focus on single animals, not multiple, interacting animals. Second, many available tools are not developed for novice users and require programming experience to run. Here, we unveil a computer vision pipeline called AlphaTracker, which requires minimal hardware requirements and produces reliable tracking of multiple unmarked animals. An easy-to-use user interface further enables manual inspection and curation of results. We demonstrate the practical, real-time advantages of AlphaTracker through the study of multiple, socially-interacting mice.

## Introduction

Behavioral analysis is considered an important core of neuroscience and research in other disciplines. The study of behaviors can be dated back to the 19^th^ century when most researchers focused on observing natural behaviors^5^. While reductionist behavioral paradigms are still widely used to study specific aspects of behavior in a controlled manner, allowing animals to freely explore more spaces and to exhibit complex behaviors greatly extends our understanding of systems neuroscience^6^.Yet, ethological behavioral research challenges our bandwidth to acquire sufficient labeled behavior data for drawing statically meaningful conclusions^7^. In addition, traditional human observations suffer from an intrinsic subjectivity and are both time and labor-intensive.

The intrinsic capacity of computational analysis for high-throughput assessments has greatly facilitated the advancement of behavioral neuroscience^8–10^. The standard workflow of behavioral analyses consists of two components in series: the tracking of the animal and behavior identification. The tracking component has benefited greatly from advances in pose estimation, such as DeepLabCut, a software package that can reliably track human-defined unique keypoints^11^. A recent algorithm, Moseq, has made progress on automated behavioral identification by using a depth camera and unsupervised learning theory^12^. SimBA presents an open-source package with a graphical interface and workflow that uses pose-estimation to create supervised machine learning^13^. However, these methods have not been effective in tracking multiple identical animals. Rapid advancement in understanding the neural mechanism of social behaviors challenges us to develop algorithms that can handle a more complicated situation, such as multi-animal tracking and the subsequent social behavioral identification. Besides the current tool we are presenting, other tools for multiple animal tracking are emerging. As an example, SLEAP is a full-featured general-purpose multi-animal pose tracking framework designed for flexibility and tested on a diverse array of datasets representative of common social behavioral monitoring setups^14^.

Here we present a behavioral analysis pipeline that allows for reliable tracking of multiple identical or near-identical mice, as well as behavioral classification. Our tracking algorithm achieves state-of-the-art performance in multi-animal tracking, even with different gear or backgrounds. Moreover, a user-friendly interface (UI) further streamlines the error handling procedure in tracking. For the behavioral identification part, we apply unsupervised machine learning for both individual as well as social behavioral classification. Finally, we provide an intuitive UI for easy inspection and modification of clustering results that does not require prior programming experience.

## Results

### Section 1: AlphaTracker

The AlphaTracker pipeline consists of three consecutive stages: tracking (AlphaTracker), behavioral clustering, and result analysis with customized UI. AlphaTracker enables multi-animal tracking on videos recorded via webcam, rendering this tool convenient and affordable for the laboratory setting. The unsupervised behavioral clustering allows for unbiased identification of behavioral motifs, the results of which can be further inspected and error-corrected in a customized user interface.

The tracking component of the pipeline (AlphaTracker) is adapted from AlphaPose^15^, a human pose estimation algorithm that benchmarks the accuracy and speed of pose estimation. The algorithm consists of three steps: animal detection, keypoint estimation and identity (ID) tracking across frames (Figure 1).

**Figure 1:**
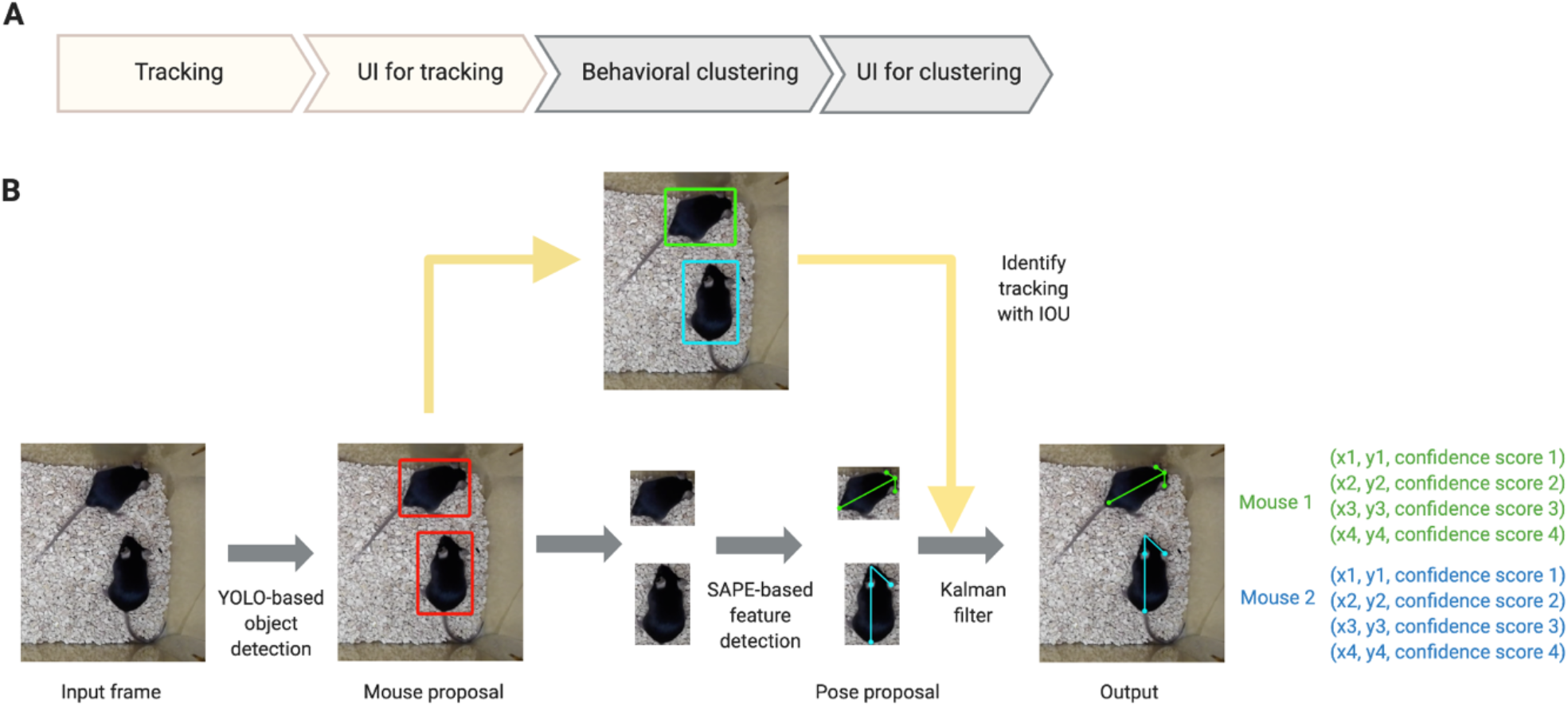
AlphaTracker architecture and pipeline. **A**. Overview of the AlphaTracker pipeline. **B**. architecture of AlphaTracker which combines object detection using You Only Look Once (YOLO) neural network, pose estimation using Single Animal Pose Estimation (SAPE), and identity tracking based on intersection over union (IOU) for subsequent frames with an additional Kalman filter to correct tracking errors.

**Figure 2:**
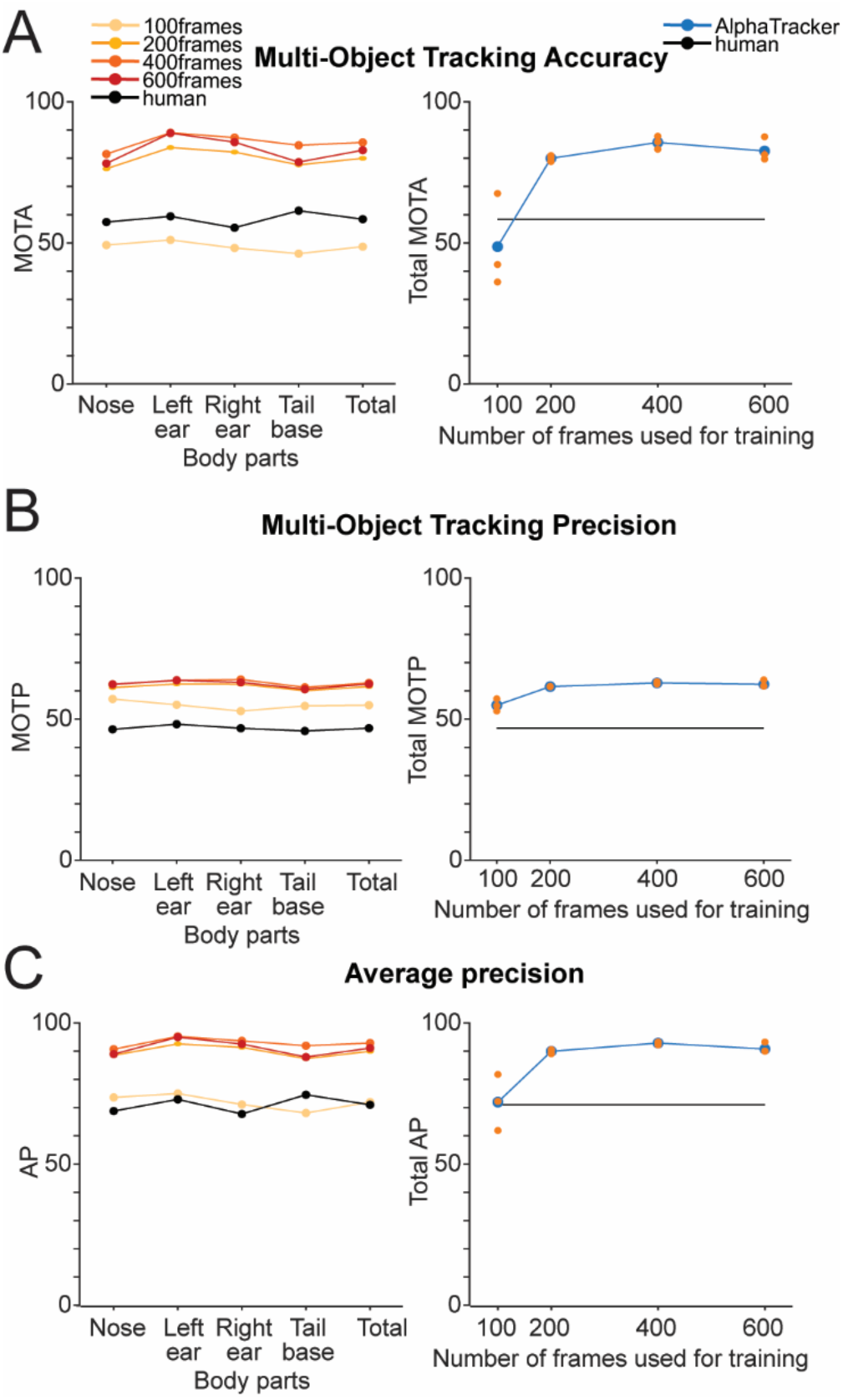
AlphaTracker is capable of tracking multiple animals with better accuracy and precision than humans. AlphaTracker (**A**) Multiple Object Tracking Accuracy (MOTA), (**B**) Multiple Object Tracking Precision (MOTP), and (**C**) Average Precision (AP) for held-out data compared to human accuracy and precision when tracking two unmarked identical mice. On the left, metrics are reported for each body part individually and on the right, the metrics are averaged across body parts and individual replicates of the testing are reported for different numbers of frames used during training.

First, the algorithm detects the positions of animals in each frame with YOLOv3^16^ which is a state-of-the-art convolutional neural network designed to detect objects at a high inference speed.

Next, individual animals are cropped out with the bounding box output from YOLOv3. The cropped individual images are fed into Squeeze-and-Excitation Networks (SENet)^17^ which estimates keypoint positions. For our mouse dataset, we chose the nose, tail-body junction and two ears as our four keypoints. The outputs from SENet include x and y coordinates as well as a confidence score which indicates the reliability of each identified keypoint. The confidence score can facilitate the identification of error-prone regions in the tracking UI.

The final step of mouse tracking is consistent tracking of ID across frames. This presents a significant challenge for many platforms as animals of the same genetic lines often look alike. Traditional Re-ID methods previously implemented^18–20^ tend to fail since such methods typically rely on differences in the appearance of the animals tracked. In our pipeline, we propose a novel target association method that can capture hierarchical visual information to tackle the challenge of keeping track of identities of nearly identical animals across frames. First, we define a descriptor for the position and orientation of the animal from the set of bounding boxes around the whole animal and each key body part. Then, we calculate the similarity of all descriptor pairs for all pairs of adjacent frames according to formula 1.

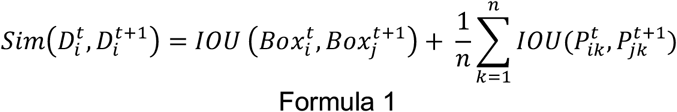

In formula 1, 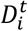 is the descriptor of animal *i* at frame *t*, 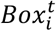 is the bounding box of animal *i* at frame *t* predicted by the convolutional neural network. 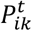 is the box that wraps the *k* th body point of animal *i* at frame *t*. Intersection Overlap Union *(IOU)* is defined by formula 2.

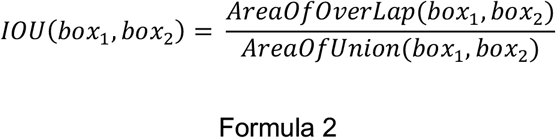

After sorting the descriptor similarities in the descending order, the descriptors between two adjacent frames with the highest similarity are matched and assigned with the same tracking ID. Then, for the dyad across frames, we match descriptors with the second-highest similarity score. This procedure is repeated until all animals in the current frame are assigned with a tracking ID or until no animals are left unmatched.

In some cases, the predictions of bounding boxes and body points may not be accurate due to either tracking errors or occlusion. To address these challenges, we adopted Kalman filtering^21^. Constructed for each keypoint to model the motion characterized by velocity and acceleration, Kalman filtering enables the prediction of keypoint locations in consecutive frames when the users amend the tracking results in the UI.

### Section 2: Performance

AlphaTracker shows reliable performance in tracking multiple unmarked, identical animals. Here, we test its performance in two popular settings in social neuroscience research.

Since environmental context plays a major role in shaping behaviors, we argue that it is highly valuable for any tracking algorithm to perform robustly across environmental settings. As opposed to the simplest setting of white unhindered open field, two more complex arenas serve as our backgrounds for training and testing: 1) home cage with bedding and 2) metal operant chambers where conditional behavior is most commonly studied. AlphaTracker achieves high accuracy in both cases (Supplemental Videos 1-2).

Another challenge in behavioral experiments derives from occlusion by head implants such as optical fibers for optogenetics and electrodes for electrophysiology analysis. These devices change the overall appearance of animals and may occlude keypoints from sight. We show that AlphaTracker performs robustly in these cases. As a side note, AlphaTracker shows good tolerance for low-resolution videos (eg. 576p) taken from webcams (Supplemental Video 3). This allows continuous monitoring of multiple subjects in parallel over a long period.

As animals in the wild tend to live in larger groups, the dynamics of which may be crucial for neuroscience research, we extend our methods to four mice home cage interaction. We show that AlphaTracker can consistently track four identical-looking C57/BL6 mice over time which greatly facilitates our social group dynamic research (Supplemental Video 4).

The AlphaTracker model can be trained quickly and easily. This training consists of two separate steps: 1) training the animal detector and 2) training the pose estimator. Training the animal detector relies only on the bounding box annotation, while training the pose estimator requires both bounding box annotation and body point annotation. Each image in the training dataset is automatically cropped according to the bounding box annotation. Cropped images are fed to the pose estimation network during pose estimator training. The initial weight of the convolutional neural network for animal detection is pre-trained on the ImageNet dataset and the initial weight of the convolutional neural network for body point estimation is pre-trained on a human pose dataset. Users can easily modify the training hyperparameters, such as the number of training epochs based on model performance while avoiding the pitfall of overfitting.

The significant time and hardware commitment for most deep learning-based models can be intimidating for many neuroscience labs. AlphaTracker provides a quick and easy solution. We tested training time as training tends to be the most time-consuming stage in the pipeline. On a server with NVIDIA TITAN Xp and Xeon(R) CPU E5-2630, it took about 0.2 seconds to train a batch of 10 images, and it took about 30 minutes for the pose estimator to converge with 6000 training images.

### Section 3: Clustering

The behavioral clustering component of AlphaTracker allows clustering of both individual behavior and social interaction in an unsupervised manner. This unsupervised approach holds great value as it minimizes the human bias in behavior identification.

The inputs for the clustering algorithm are features extracted from keypoints and a binary mask of the animal (Figure 4; Supplemental Videos 5-6). Hierarchical clustering is then performed on the units of 15 consecutive frames for all individual mouse behaviors. When two mice are within a given distance, we perform hierarchical clustering using features from both mice and their interaction (eg. distance from nose to nose) to obtain the social behavior clusters. Here, we included a total of 6194 clips for individual behavioral clustering and 2241 clips for social behavioral clustering collected from 6 videos. Our algorithm is able to capture the following individual behaviors: walking, digging, sniffing, rearing, turning, face grooming, and body grooming, and social behaviors: following, chasing, anogenital sniffing, face sniffing and social rearing.

**Figure 3:**
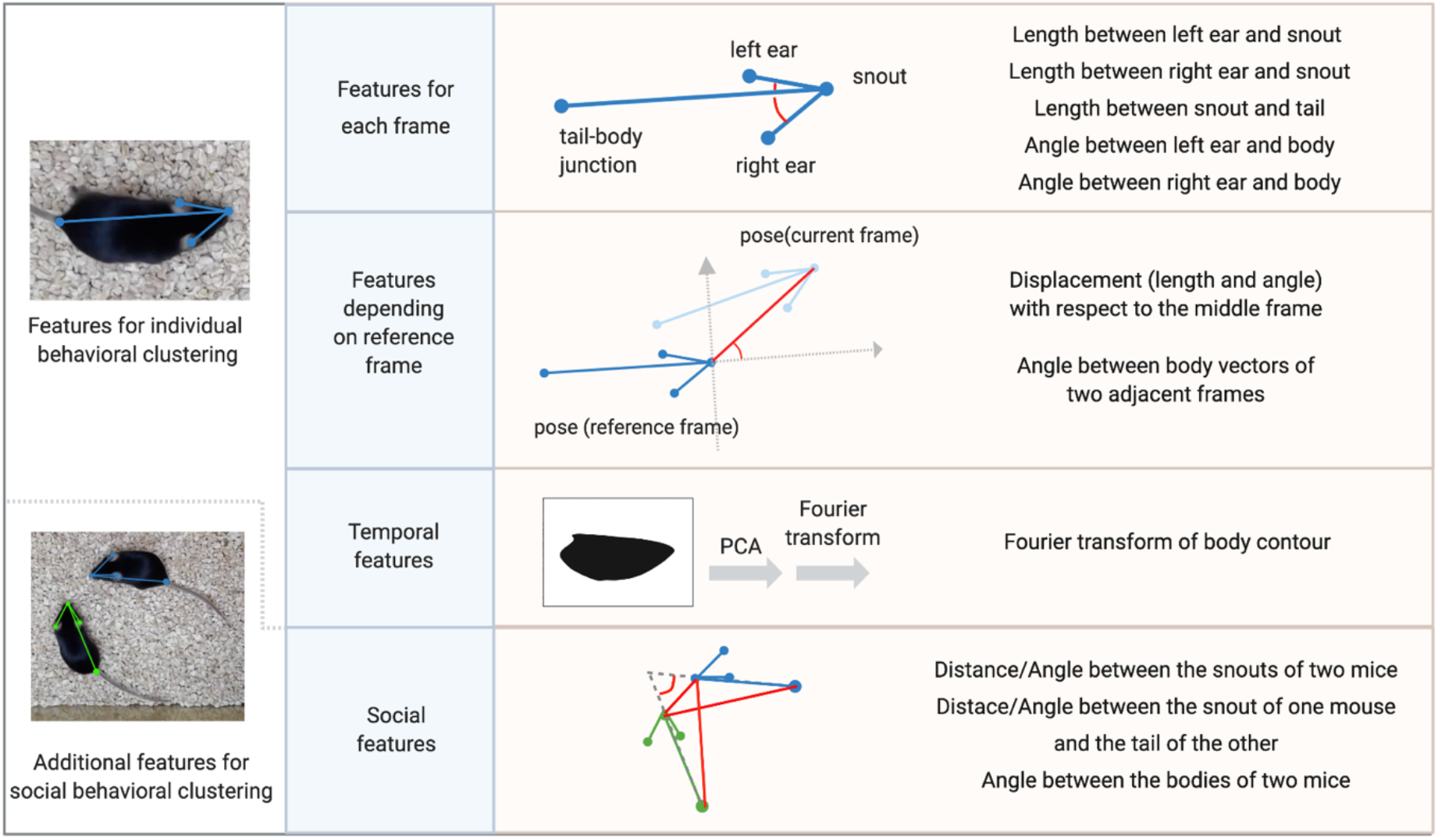
Individual behavior and/or social features are calculated based on pose estimation results. Individual behavioral clustering depends on features for each frame, features depending on the reference frame and temporal features. Social behavioral clustering depends on the individual features of both mice and additional social features. The definition of example features is depicted in the diagram. (Blue, light blue, green: pose skeleton. Red: distances between two points or angles between two vectors. Binary image: mouse contour)

**Figure 4:**
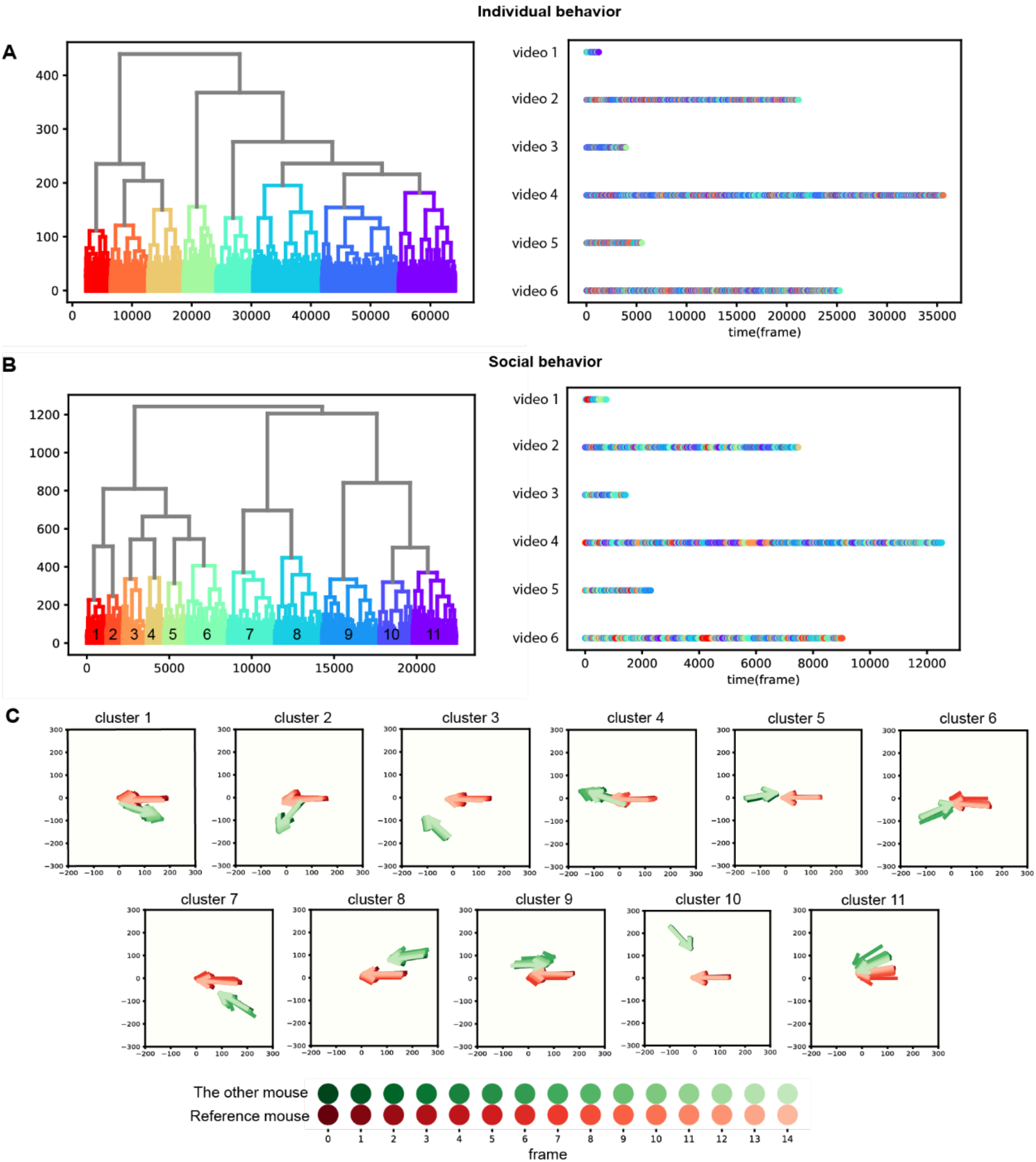
AlphaTracker enables unsupervised behavioral clustering. Hierarchical clustering was performed on clips consisting of 15 frames (500 ms duration) generated from 6 videos of interacting dyads (the 6 videos are of different durations). Dendrogram indicates the clustering results, with different clusters colored differently. Note that the colors of individual and social clustering are independent of each other. **A.** Dendrogram and timeline plot for individual behavioral clusters identified by the unsupervised clustering. The color in the timeline plot corresponds to the color in the dendrogram. **B.** Dendrogram and timeline plot for social behavioral clusters of two interacting mice. The color in the timeline plot corresponds to the color in the dendrogram. **C**. Example skeletons summarizing each social behavioral cluster. The numbering of clusters corresponds to the numbering in the dendrogram in **B**. The skeletons are aligned intentionally so that the nose of the reference mouse is at the origin and its body is on the x-axis. Colormap scale at the bottom indicates the color scheme for each frame.

**Figure 5:**
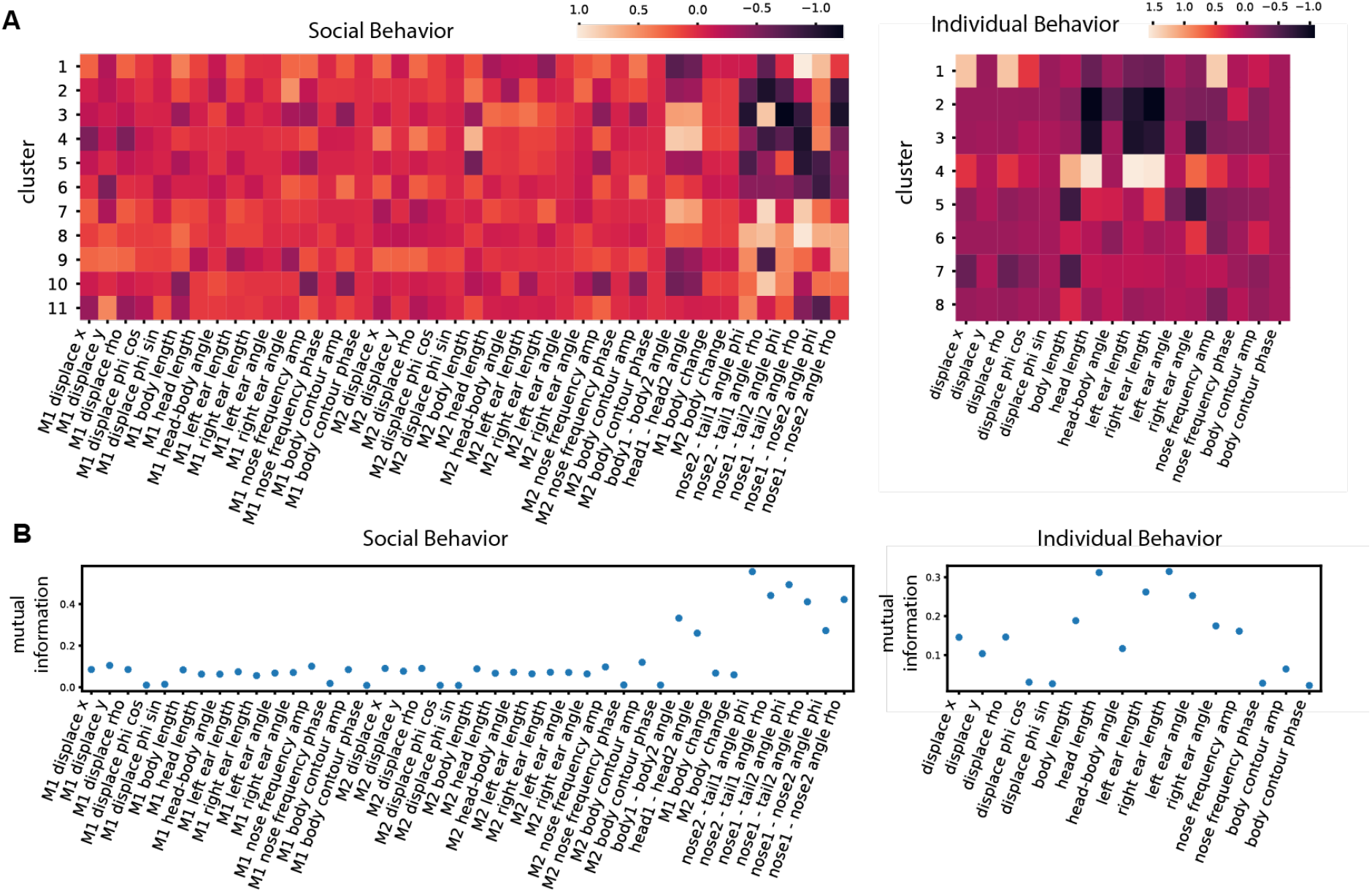
Distinct features contribute to the clustering of individual vs social behavior. **A**. Heatmap of normalized values for the different features used for clustering for each of the behavior clusters obtained with individual or social behavior data. M1 denotes mouse 1 and M2 denotes mouse 2. **B**. Mutual information between cluster assignment and each feature for social/individual behavioral clusters. A higher mutual information score indicates the feature being a salient discerning factor for that cluster.

To evaluate the importance of each feature in clustering, we calculate the mutual information between features and cluster assignment, with the expectation that higher mutual information indicates that the feature may be a unique characterization of that cluster. For example, distances between two mice are a stronger indicator in social clusters. Features related to the head such as head length and nose-left ear distance stand out among other individual features. This indicates that the head represents a salient component of many behaviors like rearing, digging and turning.

The identified behavioral clusters allow users to visualize the temporal dynamics of animal behavior. This opens up the opportunity for associative analysis between changes in behavior motifs with experimental factors like optogenetic stimulus, drug administration, environmental changes and manipulation in the social hierarchy. One limitation of this unsupervised clustering algorithm is that it requires a uniform behavioral motif length when evaluating the data.

The performance of our behavioral clustering has been validated by comparing the algorithm outputs to the ground truth of human annotation. A human scorer was asked to annotate behaviors into different categories. We measured the similarity of the class assignment between the algorithm and human scorer with Adjusted Rand Index (ARI). Independent labels are expected to have a negative *AR*I, clustering labels that are close to the ground truth labels would have a positive *ARI* and random label assignments would have an *ARI* score close to zero.

We compared the performance of the algorithm on datasets with different sizes. The small dataset contains five videos including two videos with human-annotated ground truth (1345 clips in total), while the large dataset contains two more videos (3034 clips in total). The results shown in Table 1 suggest that both datasets perform significantly better as compared to randomly assigning clips to each cluster. Moreover, the performance of the model further improved when given a larger clustering dataset, likely due to better coverage of the continuous input space. *ARI* also serves as a useful metric for model performance when users were tuning parameters for clustering.

**Table 1.**

|  | Large dataset<br>(3034 clips) | Small dataset<br>(1345 clips) | Random<br>assignment |
| --- | --- | --- | --- |
| Adjusted rand index | 0.201186 | 0.186533 | 0.003451 |

AlphaTracker was developed and validated on Linux environments with GPUs installed. If users do not have access to Linux or GPUs, AlphaTracker can also be operated on Google Colaboratory (Colab). Colab provides ~12 hours of uninterrupted access to GPUs which is sufficient for the training and testing of AlphaTracker. Results can be downloaded from Colab and fed into the user interfaces that run locally. The time required for training and testing on Colab is roughly equal to those on local computers with GPUs. A guide to using AlphaTracker on Colab can be found in our Github repository.

### Section 4: User Interface

Given that no deep learning framework is error-free, we designed customized user interfaces (UIs) for inspecting and revising both the tracking and clustering results (Figure 6; Supplementary video 7). Our UIs are built upon a web-based framework that enables the user to run this application on any system with a pre-installed Python environment.

**Figure 6:**
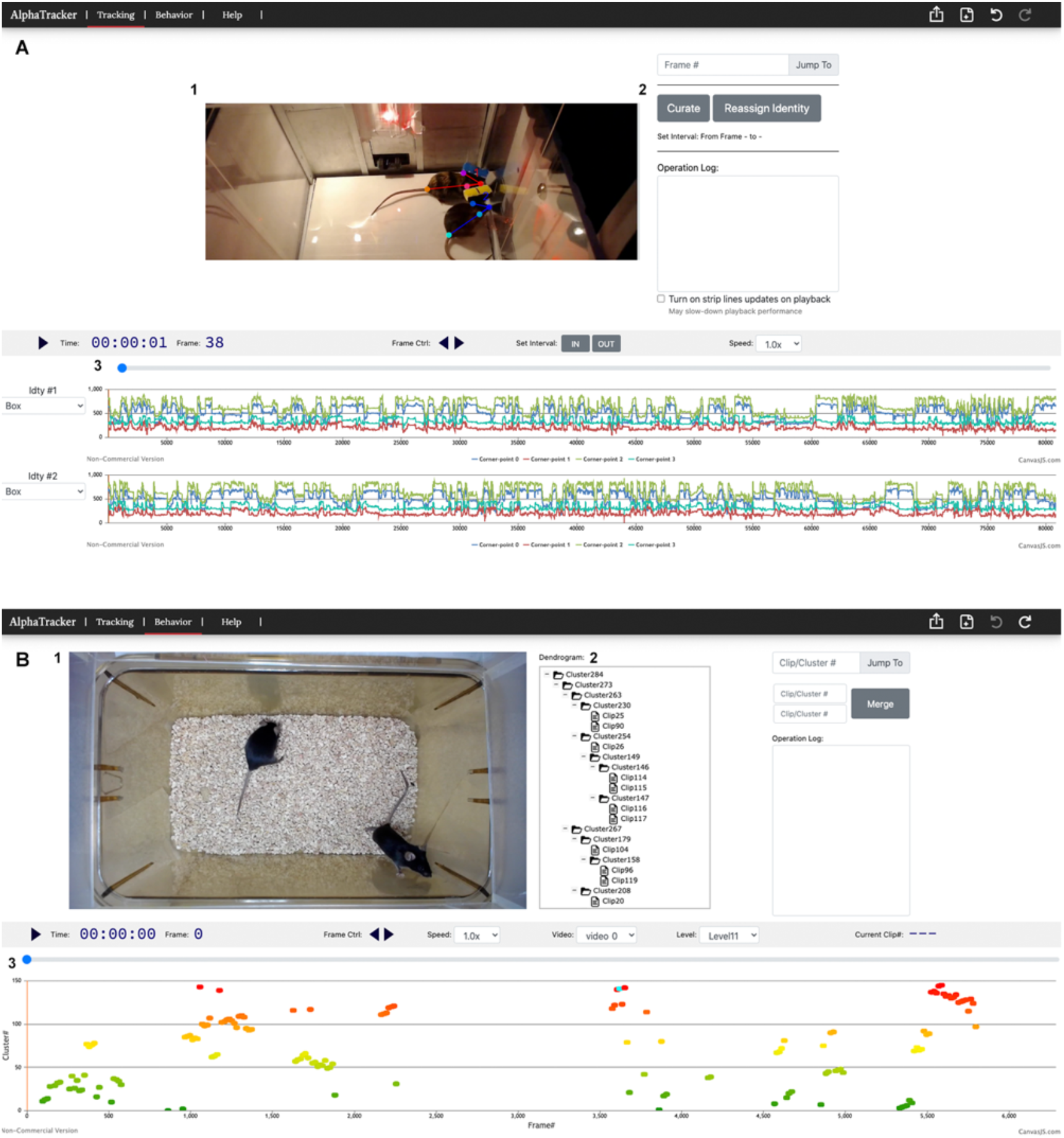
User Interface for AlphaTracker provides a solution for easily fixing errors and inspecting results. **A**. UI for tracking: Area 1 displays the video with the keypoints tracked and allows users to interact with and modify these points. In Area 2, users can re-assign identities and curate the keypoint sequences. Area 3 enables the visualization of keypoints and confidence levels over time. **B**. UI for behavioral clustering: Area 1 displays the imported video used for clustering. Area 2 shows the dendrogram of the clustering results. Users can modify the clustering results by moving, merging branches within the dendrogram. Area 3 is the visualization of clip IDs over time. Each dot is a clip and users can toggle over the clips to see their cluster assignment.

The tracking UI allows the user to scroll through the video and to inspect whether keypoints are correctly positioned, and whether mouse IDs are correctly tracked across time. The event timeline enables the user to zoom in and out to reveal more/fewer details. To facilitate error identification, the confidence score outputted by the tracking algorithm is plotted on the timeline to suggest potential periods of low confidence levels. The confidence score plot can be toggled on/off when needed.

The tracking UI also enables easy correction of errors. For keypoints that are wrongly positioned, users can drag the keypoint to the correct position. To correct flipped identities, users can reassign their identities by clicking the “reassign” button. While the above-mentioned corrections are performed on one frame, the subsequent frames are likely to suffer from the same errors. To automatically correct such errors, we introduce a function called “curate”. Users can select a range for the adjustment to take effect, then click the “curate” button which automatically computes the adjustment for subsequent frames. Results from the tracking UI consist of keypoint coordinates for each mouse and are stored in a JSON file format in a similar fashion to the data generated by the tracking algorithm.

The clustering UI features an interactive dendrogram which displays the clustering results. By default, the dendrogram is fully expanded to the lowest level. If users wish to hide the details and view only the broader cluster assignment, one can set the dendrogram to expand to a specified level. The scatterplot in the timeline allows users to inspect the distribution of clips belonging to certain clusters. Upon selection of a cluster in the dendrogram, the corresponding dots representing clips of that cluster will be highlighted in cyan in the timeline. Users can also navigate to a clip of interest by putting its index in the navigation textbox.

The clustering UI also enables convenient modification of the clustering results. Our UI allows easy renaming of the clusters such that users can name motifs in ways that describe the behavior. To change the cluster assignment, users simply move clusters/clips to other clusters. They can also merge clusters of the same level to reduce redundancy. The curated clustering results are saved in a JSON file for subsequent analysis.

### Conclusion

In summary, we provide a behavioral analysis platform called AlphaTracker which incorporates multi-animal tracking, pose estimation and unsupervised behavioral clustering to facilitate ethology and social behavioral research. Our tracking algorithm achieves the state-of-art accuracy of multi-animal tracking which lays the foundation for stringent biological studies. This analysis pipeline can be readily adopted by most research groups given its minimal requirement for hardware and efficient training procedure. Furthermore, the customized UI further allows easy inspection and modification of clustering results without the need for additional programming. We hope that AlphaTracker becomes a useful platform for the community of researchers interested in social behaviors and that it facilitates novel biological insights.

## Methods

### Dataset

The training dataset for the animal detection and body points estimation neural network consists of frames randomly selected from the videos to be analyzed. To reduce the sample size for labeled frames, one can start with annotating a few hundreds of randomly selected frames and annotate more if needed. Labeled frames are automatically split and used for training and validation. Our training dataset represents frames from videos of mouse dyads interacting in their home cages.

### Metrics for tracking

800 distinct frames were randomly sampled from 4 different videos of the resolution (1920×1080p) with similar backgrounds (200 frames per video). The bounding box, nose, left ear, right ear and tail base of each mouse in those frames were labeled manually. Human annotators also kept track of the identities via consistently following the same mouse during annotation. To ensure a fair comparison, human annotators took note of whether the body parts were occluded.

To evaluate our multi-animal tracking model, we used the standard metrics in multiple object tracking, the CLEAR MOT metrics Average Precision(AP), Multiple Object Tracking Accuracy (MOTA) and Multiple Object Tracking Precision (MOTP)^22^. Average Precision (AP) calculates the accuracy of object detectors using the precision and recall values. Multiple Object Tracking Accuracy (MOTA) evaluates three types of errors: missed objects in a sequence, false positives, and mismatches. MOTP examines the total position error for matched object-hypothesis pairs over all frames, averaged by the total number of matches made.

The evaluation was implemented with an open-source tool Poseval^23^. To adapt the MOT metrics to evaluate mouse tracking, we modified the threshold of distinguishing matched keypoints from mismatched keypoints to be 5% of the diagonal of the bounding box. We trained our model with different split ratios between the training and evaluation datasets (75%, 50%, 25% and 12.5%, 3 replicates for each ratio). We made sure there was no overlap between the training and evaluation datasets. For example, for the split ratio of 75%, we randomly selected 150 consecutive frames from each of the four videos to constitute a total of 600 frames to train the network.

To compare the performance of our tracker to human performance, we also included the CLEAR metrics of human annotation by comparing the annotation of 200 frames done by 2 different people.

### Clustering algorithm

AlphaTracker uses hierarchical clustering^24^ to classify mouse behavior. The rationale for using hierarchical clustering includes: 1) Animal behavioral taxonomy is intrinsically hierarchical in structure. 2) It allows intuitive re-organization of the results once the linkage matrix is computed. Users can change the cluster assignment of clips and merge/delete clusters with the UIs we provide.

Each video is divided into video clips of the same length defined by users. The video clip length chosen in this paper was 500 milliseconds (15 frames). Features are calculated for individual mice in each clip. For social behavioral clustering, we also calculate social features based on the pose of two mice.

Agglomerative hierarchical clustering algorithm^25^ is applied to the calculated features. For a dataset with n clips, hierarchical clustering starts with n clusters, with each cluster consisting of one clip. During each iteration, the distance between each pair of clusters is computed. Two clusters with the shortest distance are removed from the cluster set and combined to form a new cluster. The algorithm stops when only one cluster remains. The final result of the clustering is represented as a dendrogram with each leaf representing one clip. The two legs of the U-link indicate which clusters were merged. The length of the two legs of the U-link represents the distance between the child clusters^25^.

We compute the distance^26^ between cluster u and cluster v with formula 3

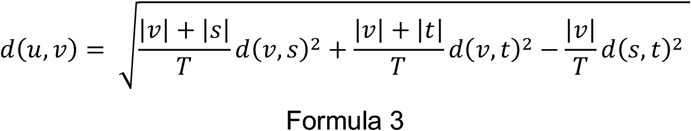

where u is the newly joined cluster consisting of clusters s and cluster t, v is an unused cluster in the cluster set. |*| is the cardinality of the argument. *T* = |*v*| + |*s*| + |*t*|. The initial distances between the clusters are calculated with Squared Euclidean distance. Note that we only need to calculate the distance between the newly joined cluster and the other unused clusters at each iteration, since the distances between the other clusters remain the same.

### Features

To conduct hierarchical clustering on mouse behaviors, we calculate features based on the estimated poses: nose, left eye, right eye and body-tail junction. To obtain a coherent feature representation for different video clips, we normalize the features before clustering. For each video, users first choose the reference mouse. We apply a coordinate transformation so that the nose of the reference mouse in the middle frame of the clip is set as the origin, and the x-axis aligns with the reference mouse’s body vector in the middle frame of the clip. After computing the following features, users can assign different weights to each feature to reflect their relevant importance in behavioral clustering. The features we extract from the clips are as follows:

#### Features for individual behavior

- The head vector points from the nose to the midpoint of the line segment connecting the left ear and the right ear. Head length is defined as the norm of the head vector (‘head length’).
- The body vector points from the nose to the tail base. The body length is defined as the norm of the body vector (‘body length’).
- The ear vector points from the nose to the ear. The ear-nose length is defined as the norm of this ear vector (‘left ear length’, ‘right ear length’) The angles between ear vectors and the body vectors are also included in the features (‘left ear angle’, ‘right ear angle’)
- The angle between the head vector and the body vector is included (‘head-body angle’).
- The displacement vector of the nose is included (‘displace x’, ‘displace y’, ‘displace rho’, ‘displace phi cos’, ‘displace phi sin’).
- The angle between the body vector and that of the previous frame is included (body change’).
- To emphasize the temporal dynamics of displacement, we apply the Fourier transformation to the displacement of noses between two frames (‘nose frequency amp’, ‘nose frequency phase’).
- To capture the information in the soft body shape, the binary mask of each mouse is extracted, followed by dimension reduction with Principal Component Analysis (PCA). The cut-off threshold for eigenvalues is chosen to retain 90% of the variation. We then apply the Fourier transformation to capture the temporal features of contour information (‘body contour amp’, ‘body contour phase’).

#### Features for social behavior

- Angles between the body vectors of two mice (‘body1 - body2 angle’)
- Angles between the head vectors of two mice (‘head1 - head2 angle’)
- Distances and angles from one mouse’s nose to the other mouse’s tail (‘nose2 - tail1 angle phi’, ‘nose2 - tail1 angle rho’, ‘nose1 - tail2 angle phi’, ‘nose1 - tail2 angle rho’, ‘nose1 - nose2 angle phi’, ‘nose1 - nose2 angle rho’)

### Demo Skeletons for each cluster

Selected features were averaged across clips of the same cluster to form the representative feature vector for the cluster, including the body length, the left ear-nose length, the right ear-nose length, the distance between the noses of two animals and several related angles. These features allowed the reconstruction of demo skeletons representing each cluster. For easy comparison between clusters, we set the nose of the reference mouse as the origin and its body vector as the x-axis.

### Plots for clustering data

The dendrogram was generated based on the linkage matrix produced by the clustering algorithm. Branches below the user-defined threshold were colored according to their cluster assignment. The threshold is set to be 500 for individual behavior and 200 for social behavior. The timeline plots showed the cluster assignments for each clip (15 frames in length), with their color matching the cluster assignment in the dendrogram. The heatmaps were generated with Scipy^25^ after normalizing each feature to have a mean of zero and a standard deviation of 1. Mutual information was calculated using the *mutual_info_classif* function from Scikit-learn package^27^ which quantifies the mutual information between each feature and the cluster assignment.

### Evaluation metrics—Adjusted Rand index

Suppose *C* represents the labels given by human annotation and *K* represents the label given by hierarchical clustering. We first calculate a useful metrics Rand index (RI):

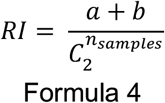

where *a* is the number of clip pairs that are grouped in both *C* and *K, b* is the number of clip pairs that are in different sets in both *C* and *K*, 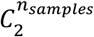 is the total number of possible pairs. Adjusted Rand Index(*ARI*) is then calculated:

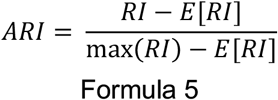

where *E*[*RI*] is the expected *RI* for random labels.

### User Interface

To ensure the compatibility of the UI with most platforms, we utilize web-based frameworks like Bootstrap and jQuery. The tracking UI is powered by JavaScript, with the “curate” function being dependent on Python. The use of PyIodine library enables the app to run Python codes and integrate JavaScript variables. For data visualization, CanvasJS is adopted for its superior performance and user-friendly functionality (eg. zoom-in or zoom-out). This library is also deployed in the clustering UI to visualize cluster distribution in the timeline.

The AlphaTracker code is available here: https://github.com/ZexinChen/AlphaTracker

## Notes

https://github.com/ZexinChen/AlphaTracker

### Competing Interest Statement

The authors have declared no competing interest.

